# The conserved biochemical activity and function of an early metazoan phosphatidylinositol 5 phosphate 4-kinase regulates growth and development

**DOI:** 10.1101/2024.08.01.606031

**Authors:** Harini Krishnan, Suhail Muzaffar, Sanjeev Sharma, Visvanathan Ramya, Avishek Ghosh, Ramanathan Sowdhamini, Padinjat Raghu

## Abstract

The ability to co-ordinate function between multiple cells is a critical requirement for multi-cellularity. This co-ordination is mediated by hormones or growth factors, molecules secreted by one cell type that can convey information to the other cells and influence their behaviour. Hormone-dependent signalling is mediated by second messenger systems;phosphoinositides (PIs) generated by lipid kinase activity are one such key second messenger system. Phosphatidylinositol 5 phosphate 4-kinase (PIP4K) is a lipid kinase that phosphorylates phosphatidylinositol 5-phosphate (PI5P) to generate phosphatidylinositol 4,5 bisphosphate [PI(4,5)P_2_]. Following a comprehensive bioinformatics analysis of ca. 23296 proteomes covering the tree of life, we find that PIP4K is a metazoan-specific enzyme, although its homologs are also found in choanoflagellate genomes. To understand their function in early metazoans, we experimentally analysed the biochemical activity and physiological function of PIP4K from several early metazoans. We find that the PIP4K enzyme from an early branching metazoan sponge *Amphimedon queenslandica* (AqPIP4K), regarded as the earliest evolved metazoan, shows a biochemical activity highly conserved with human PIP4K; AqPIP4K is able to selectively phosphorylate PI5P to generate PI(4,5)P_2_ just as effectively as the human enzyme. Further, AqPIP4K was able to rescue the reduced cell size, growth and development phenotype in larvae of a null mutant in *Drosophila* PIP4K. These phenotypes are regulated through activity of the hormone insulin, acting via the cell surface insulin receptor, a member of the receptor tyrosine kinase family, that is unique to metazoans. Together, our findings indicate that in early metazoans, AqPIP4K is likely to function in a signal transduction pathway that is required for receptor tyrosine kinase signalling. Overall, our work defines PIP4K as a signal transduction motif required to regulate receptor tyrosine kinase signalling for intercellular communication in the earliest forms of metazoa.

## Introduction

A key requirement for the successful survival of multi-cellular organisms is the ability of its constituent cells to communicate. This process of communication, referred to as signalling is mediated by molecules in the extracellular space that convey information to individual cells; for example, hormones such as insulin, *wnt* ligands and TGFβ are small molecules that aid communication between cells. Once signalling molecules are released, they bind to ligand-specific receptors leading to biochemical reactions that trigger intracellular signalling, convey information, and lead to changes in cell state or behaviour. In line with these requirements, genes encoding such ligands and their cognate receptors have been identified in the genomes of the first evolved multi-cellular organisms ^1,2^ and the evolution of such signalling systems must be a key requirement for multicellularity.

Among the intracellular signal transduction systems used by eukaryotic cells are the phosphoinositides (PI). PIs are glycerophospholipids localized to the cytoplasmic surface of organelle membranes. They can be generated in response to extracellular stimuli and serve as signalling molecules for intracellular communication ^3,4^. For example, the binding of insulin to the insulin receptor (InR), a receptor tyrosine kinase, triggers the activation of Class I phosphoinositide 3 kinase (PI3KC1) leading to production of phosphatidylinositol 3,4,5 trisphosphate (PIP_3_). PIP_3_ acts as an intracellular signalling molecule regulating processes such as cell size, cell division, adhesion and motility. Although most components of the PI signalling cascade are conserved in all eukaryotes, PI3KC1 and PTEN, the PIP_3_ phosphatase that dephosphorylates PIP_3_ are found only in metazoan genomes ^5,6^, suggesting a specific role for PIP_3_ dependent processes in metazoans.

Given the uniqueness of PIP_3_ signalling to multicellular organisms, regulators of PIP_3_ production and signalling might also be unique to metazoans. For e.g. receptor tyrosine kinases (e.g EGFR, InR, etc) whose activation leads to PI3KC1 activation are reported only in metazoan genomes ^7, 8^. A more recently described regulator of PI3KC1 signalling is phosphatidylinositol 5 phosphate 4-kinase (PIP4K). PIP4K is an enzyme that phosphorylates phosphatidylinositol 5-phosphate (PI5P) to generate phosphatidylinositol 4,5 bisphosphate [PI,(4,5)P_2_] (**Figure 1A**) ^9,10^. Genes encoding PIP4K are not found in the genome of the unicellular yeast *S. cerevisiae* but 3 genes encode PIP4K in mammalian genomes^11^. Further, in *Drosophila,* that contains only a single PIP4K (*dPIP4K*), this protein appears to regulate insulin signalling^12^ and recent studies in *Drosophila*^13^ and human cells ^14^ have shown that PIP4K is a key regulator of PI3KC1 signalling. However, the evolution and function of PIP4K with respect to the tree of life remains unclear. Here we report that PIP4K genes are not found in unicellular eukaryotes but are found in all metazoan genomes. Importantly PIP4K is found in the genomes of the earliest examples of metazoans including the demosponge *Amphimedon queenslandica* (AqPIP4K) as well as in the genomes of choanoflagellates^15^. We present data that *in vitro*, AqPIP4K is catalytically active and displays the characteristic substrate specificity of mammalian PIP4K. Remarkably, AqPIP4K appears functionally equivalent *in vivo*, is able to regulate insulin activated PI3KC1 signalling and restore function in null mutants of *Drosophila* PIP4K. Together, these findings identify AqPIP4K as an ancient and evolutionarily conserved lipid kinase that regulates growth in multi-cellular organisms.

**Figure 1:**
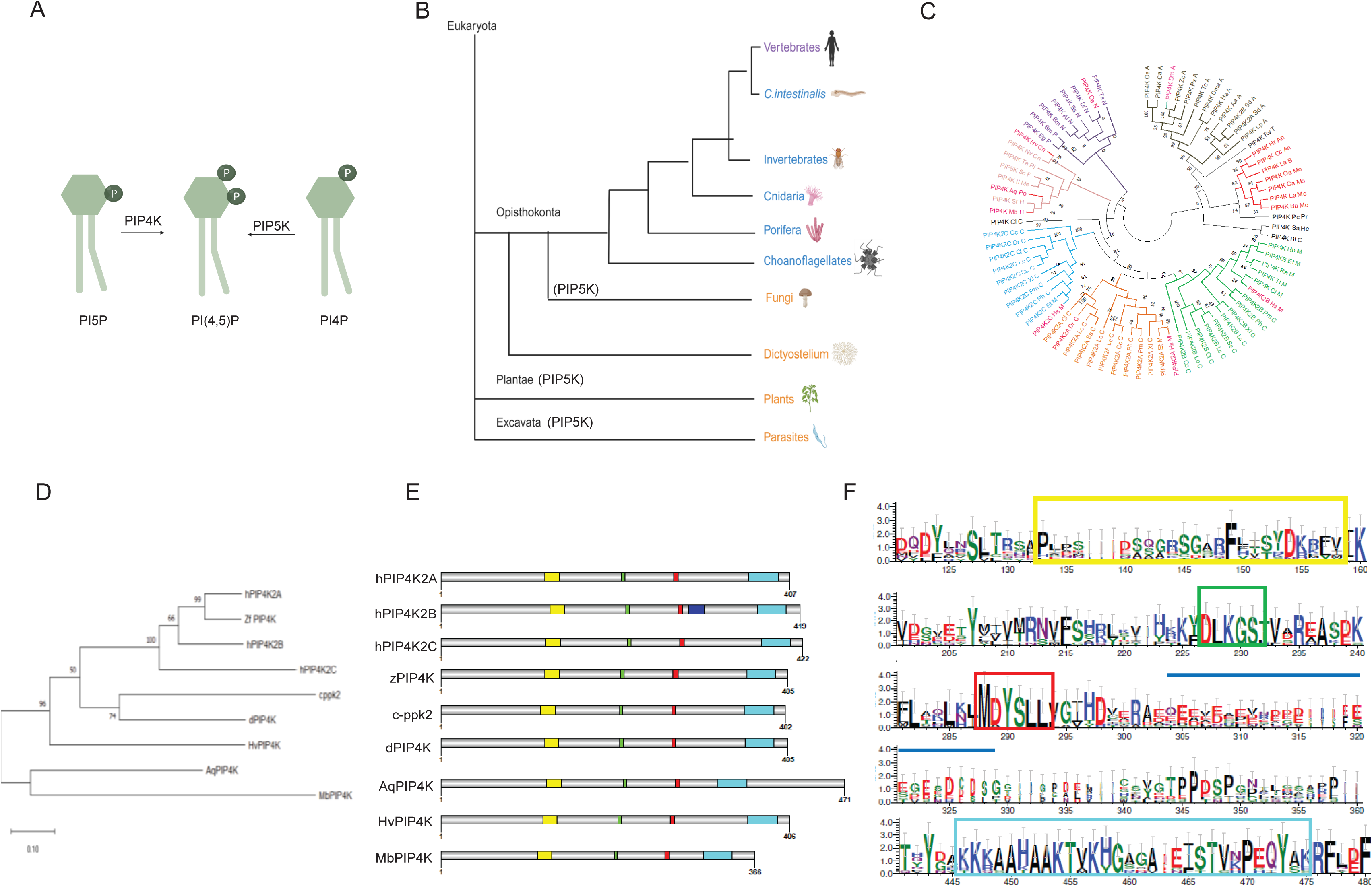
**(A)** Enzyme reaction for PIP4K and PIP5K. PIP4K converts PI5P to PI(4,5)P_2_ while PIP5K converts PI4P to PI(4,5)P_2_. **(B)** Simplified tree of life diagram for eukaryote evolution to explain evolution of PIP4K enzyme. All the groups express PIP5K. The groups marked in orange do not encode the PIP4K gene in their genome. The groups marked in blue contain a single copy of PIP4K gene in their genome while the group marked in violet contain more than one copy of the PIP4K gene. **(C)** Phylogenetic analysis of 77 PIP4K sequences to check the distribution of PIP4K sequences of organisms from different phylum. Each phylum is distinguished from other by way of difference in colour. The sub clusters have been numbered (1- early metazoans, 2- Nematodes, 3- Arthropods, 4- Molluscs, 5- Vertebrates (isoform PIP4K2C), 6- Vertebrates (isoform PIP4K2A), 7- Vertebrates (isoform PIP4K2B). The sequences selected for analysis have been marked in pink. **(D)** Phylogenetic analysis of selected PIP4K sequences from model organisms of interest. PIP4K from *A.queenslandica* has been marked in rectangular box. **(E)** Conserved domains and functionally important residues mapped on selected PIP4K sequences. Yellow -PI5P binding domain, Green and Red-ATP binding motifs and Cyan-Activation loop. **(F)** The conserved motifs and domain mapped at the amino-acid residue level using LOGO plot from selected PIP4K sequences. The x-axis denotes the residues number. The y-axis denotes relative conservation of the residues. Greater the value of Y-axis, higher the conservation. The amino acids are colour coded based on their chemical properties.

## Methods

### Identification of PIP4K enzyme sequences from metazoan genomes

The human PIP4K2A sequence was used as query to search for PIP4K sequences in the Uniprot-KB database (https://www.uniprot.org/help/uniprotkb) and the NCBI proteome dataset (https://www.ncbi.nlm.nih.gov/protein/). Independently, annotated phosphatidylinositol kinase and phosphatidylinositol phosphate kinase protein sequences were selected as start point sequences (query sequences) to search the available proteome data on the NCBI sequence database. The protein sequences for phosphoinositide kinase enzymes used for this search are listed in **Supplementary Table 1** and were obtained from NCBI sequence database. From the hits obtained by these methods, based on the presence of only a single PIPK domain and the characteristic conserved alanine residue in the activation loop, all PIP4K sequences were selected. The sequence IDs of the selected PIP4K sequences were used to map the taxonomy of the organism from which the sequence originated using the UniProt-taxonomy search tool. Based on availability and completeness of whole proteome data, one representative organism from each phylum was used for further analysis **(Supplementary Table 2)**.

The selected proteomes (**Supplementary Table 2**) were converted into a database using the makeblastdb option of Blast tool ^16^. This database was queried for PIK and PIPK sequences in each of the organisms using the query sequences (**Supplementary Table 1**). The proteome of individual organisms was searched using each enzyme from the query database at an e-value of 10^-5^ and query coverage of 70%. These parameters reduce the possibility of obtaining false positives. The length of the protein sequence of each of the query enzymes is known. Thus, out of the hits obtained from BlastP, the hit that matched the known protein length and showed domain conservation were selected further. The presence or absence of all known PI and PIP kinases were checked for each proteome.

### Identification of functionally conserved motifs and domains

The sequences annotated as PIPK domain encoding enzymes from each phylum were aligned using ClustalO^17^ and domains were mapped using Pfam annotations. The lipid binding motifs and ATP binding residues were marked from data obtained from earlier studies on these kinases. Based on the residue difference between PIP5K and PIP4K in the activation loop^18^ PIP4K sequences were identified. The PIP5K and PIP4K sequences identified in each of these organisms has been listed in **Table 1**. The hits obtained from other query sequences (PI3K and PI4K) were used to verify the presence of annotated lipid kinases in the selected organisms, especially in genomes where PIP4K sequence was not identified.

**Table 1:** List of PIP5K and PIP4K sequences identified from selected metazoan genomes.

| **Name of the organism** | **Representative phylum** | **PIP5K sequence ID** | **PIP4K sequence ID** |
| --- | --- | --- | --- |
| *Dictyostelium* | Amoebozoa | XM_003292336.1 | - |
| *Arabidopsis* | Plantae | XM_012625114.1 | - |
| *Neurospora* | Fungi | XM_009855165.1 | - |
| *Saccharomyces* | Yeast | KZV12447.1 | - |
| *Monosiga* | Holozoa | XM_001745768.1 | XM_001745565.1 |
| *Amphimedon* | Porifera | XM_001745768.1 | XM_003383668.2 |
| *Nematostella* | Cnidaria | XM_001633017.1 | XM_001647481.1 |
| *Helobdella* | Annelida | XM_009028061.1 | XM_009033250.1 |
| *Lottia* | Mollusca | XM_009061272.1 | XM_009068559.1 |
| *Drosophila* | Insecta | ABI31111.4 | NM_001038716.2 |
| *Caenorhabditis* | Nematoda | NM_05917.6 | NM_065099.4 |
| *Branchiostoma* | Chordata | XM_002591315.1 | XM_002599441.1 |
| *Homo sapiens* | Chordata | NM_001135638.2/<br>NM_003558.4/NM_012398.3 | XM_002717397.2/<br>NM_003559.4/NM_024779.4 |

### Phylogenetic conservation of PIP4Ks

PIP4K protein sequences were obtained from NCBI proteome resource as detailed above. 1046 sequences from all kingdoms of living organisms were selected. 100% redundancy cut-off was used to further remove similar sequences. In cases where sequences that belong to organisms from same phylum were more than 70% similar, only one of the sequences was selected to avoid bias in identifying conserved amino acid positions. Finally, 76 PIP4K protein sequences were selected for final analysis. MSS4 protein from *S. cerevisiae* (PIP5K in yeast) was selected to root the phylogenetic tree. MEGA 7^19^ was used to generate the phylogenetic tree.

### Identifying orthologs for genes involved in insulin signalling

The genes involved in the insulin signalling pathway that regulates growth were selected from literature analysis (insulin peptide, insulin receptor, Akt, mTOR, S6 kinase and RTK). The key, well-established components of the pathway were shortlisted and the protein sequences of human genes for these components were downloaded from NCBI. Proteomes of *Amphimedon queenslandica* and *Drosophila melanogaster* were downloaded from NCBI ftp portal. The proteomes were converted to searchable databases using makeblastdb option of BLAST tool. The homologs of human and fruit fly proteins involved in insulin signalling were identified in these proteomes using Blastp.

### Identification of all lipid kinases and phosphatases in *A.queenslandica* genome

Human and *Drosophila* protein sequences for all lipid kinases and phosphatases ^3^ were obtained from the NCBI protein sequence database. These were used as queries to search for homologs in the proteome of *A.queenslandica*. The hits obtained were classified as true homologs based on the query coverage, sequence identity with the query sequence and presence of similar domains as in the query sequence.

### Cloning of primitive PIP4Ks into a plasmid compatible for S2 cell transfection and fly transgenics generation

The cDNA from *Amphimedon queenslandica* was cloned from cDNA synthesised from a piece of sponge tissue. Primers specific to PIP4K gene at the 5’ end and 3’ end were designed and the *AqPIP4K* gene was amplified using PCR and cloned into an intermediate clone jet vector using the CloneJET PCR Cloning Kit (Cat No. K1231, Thermo Fisher). *AqPIP4K* was subsequently sub-cloned into pUAST-attB vector with a c-terminal GPF tag between the EcoRI and XhoI restriction sites. The pUAST-attB vector is compatible for gene expression in fly systems and *Drosophila* S2 cells. The pUAST-attB plasmid containing the *AqPIP4K* was used to generate transgenic flies and for transfection in S2 cells. The PIP4K from Choanoflagellate (*Monosiga breviolis,* MbPIP4K) was synthesized based on available DNA sequence information available in NCBI. In the case of Cnidaria, the *Hydra vulgaris,*(HvPIP4K) gene was cloned from a cDNA synthesis made from *Hydra vulgaris* tissue (Dr. G. Ratnaparkhi). Subsequent sub-cloning steps were performed as described for *A. queenslandica* PIP4K.

### Expression and localization of *AqPIP4K i*n S2 cells

S2 cells were cultured in Schneider’s insect medium containing 10 % FBS (Foetal bovine serum), penicillin and streptomycin. Cell were transfected using Effectene (Qiagen). After 24 hours cells were stained with DAPI (1:5000) and fixed with 4% paraformaldehyde (EMS) and imaged using confocal microscope (Olympus FV 3000) for GFP using the 60× 1.4 NA objective.

### Bacterial growth, plasmids, and cloning for experiments in Yeast

*E. coli* DH5α was used for plasmid preparation and cloning of *Amphimedon*, yeast, and human phosphatidylinositol phosphate kinases. Standard growth conditions and cloning protocols were used. 1.3 kb *Amphimedon* PIP4K (AqPIP4K) along with GFP was cloned in inducible yeast expression vector PYES2 at *EcoR1* and *XbaI* sites. MSS4 (yeast PI4P 5-kinase) was cloned in pCR-XL-TOPO plasmid and then subcloned in PYES2 plasmid at *BamHI* and *Xho*I sites..hPIP4K2A (1.3 kb cDNA) was cloned in TOPO2.1 plasmid and then subcloned in PYES2 plasmid at *BamHI* and *Xho*I sites. hPIP5K1A (1.6 kb cDNA) was cloned in TOPO-XL plasmid and then subcloned into PYES2 plamsid with a c-terminal GFP tag at EcoRI and XhoI sites.

### Yeast strains, media, and growth conditions

Background yeast strain JK9–3d and isogenic temperature-sensitive mutant SD102 (MH272–1da mss4::HIS3MX6/YCplac111::mss4–2^ts^) were kindly gifted by Prof. Michael N Hall, University of Basel, Switzerland. Complete media (YPD) and synthetic minimal media (SD) supplemented with the appropriate amino acids were used for yeast cell growth and plasmid maintenance. Standard growth conditions were used for yeast strains. Yeast cells were transformed with PYES2 plasmid carrying *Amphimedon* and human PIP4K using the lithium acetate method^20^. Expression of the AqPIP4K and MSS4::GFP fusion protein was induced by the addition of 2% galactose for 12 hours. Cell viability was determined by the number of colony-forming units of experimental and control strains at permissive and non-permissive (37^0°^C) temperatures.

### Lipid extraction

*S. cerevisiae* cells were grown at permissive temperature (28 or 30^°^C) and non-permissive temperature (37^°^C) till the mid-log phase in a shaking incubator. The culture was centrifuged at 5,000 g for 5 minutes and the pellet was washed with cold sterile water. For lipid extraction, 0.1 g of biomass was mixed with 0.2 g of glass beads in appropriate organic solvents. Cells were disrupted by agitating tubes on a vortex for 1 minutes and the process was repeated 3-5 times.

The following solvent mixtures were used for the analysis.

Lower Phase Wash Solution (LPWS) – MeOH / 1M HCl / CHCl3 in the ratio 235/245/15 (vol/vol/vol).Post derivatization Wash Solution (PDWS) – CHCl3 / MeOH / H2O in the ratio 24/12/9 (vol/vol/vol). (Note: Shake the mixture vigorously and allow settling into separate phases and use the upper phase only for washes). The supernatant was discarded and the cell pellet was resuspended in 170 µl of 1X PBS. Thereafter, 750 µl of ice-cold quench mixture was added to each tube, followed by 725 µl of CHCl_3_ and 170 µl of 2.4 M HCl. The tubes were vortexed for 2 minutes and kept at room temperature on the bench for 5 minutes and then spun at 1500 rpm in a benchtop centrifuge for 3 minutes to separate the phases. 900 µl of the lower organic phase was pipetted out into fresh tubes containing 710 µl of LPWS. The tubes were then vortexed again for 2 minutes and spun as earlier to separate the phases. The lower phase was then collected to the extent possible, without any of the upper phase, into a fresh tube.

### Lipid Derivatization

The steps until the quenching of the reaction were performed inside a chemical hood, while wearing an appropriate respirator mask. All samples were incubated on a shaker at 600 rpm for 10 minutes at room temperature after 50 µl of TMS-Diazomethane was added to each. The TMS-Diazomethane in each tube was completely quenched at the end of the 10 minutes reaction with the addition 10 µl of Glacial Acetic acid and inverting the tubes a few times and carefully snapped open immediately once after to let the Nitrogen released during quenching to escape. All sample tubes were vortexed for 2 minutes after 500 µl of the upper phase of PDWS was added to each and then spun on a benchtop centrifuge for 3 minutes to separate the phases. 400 µl of the resulting upper phase was discarded, another 500 µl of upper phase of PDWS was added to each tube and the vortex and spin steps were repeated to separate the phases. Finally, the entire upper phase was discarded from each sample. 45 µl MeOH and 5 µl H_2_O was added to each tube, mixed and spun down. Samples were then dried in a SpeedVac at 500 rpm for around 2 hours till only about 10-20 µl of solvent remained. A further 90 µl of MeOH was added to reconstitute the samples before being taken for injection and analysis. Chromatographic separation of samples (injected as duplicates) was performed on an Acquity UPLC BEH300 C4 column (100 × 1.0 mm; 1.7 µm particle size - Waters Corporation, USA) and a Waters Aquity UPLC system connected to an

### Liquid chromatography/Mass spectrometry (LC/MS)

ABSCIEX 6500 QTRAP mass spectrometer for ion detection. Solvent A (Water + 0.1% Formic Acid) and Solvent B (Acetonitrile + 0.1% Formic acid) were used with the following gradient protocol at a flow rate of 100 µl /min –

0-5 minutes: 55% Solvent A + 45% Solvent B

5-10 minutes: Solvent B increased from 45% to 100%,

10-15 minutes: Solvent B at 100%,

15-16 minutes: Solvent B reduced from 100% to 45%,

16-20 minutes: 55% Solvent A + 45% Solvent B.

We first employed Neutral Loss Scans on the mass spectrometer during pilot experiments on biological samples to search for parent ions that lose neutral fragments with masses 490 a.m.u and 382 a.m.u indicative of PIP_2_ and PIP species respectively as described in (Clark et al., 2011). Thereafter, we quantified PIP and PIP_2_ in biological samples using the selective Multiple Reaction Monitoring (MRM) method in the positive ion mode. We found lipid species with the combined 2-chain carbon lengths and double-bond numbers of 32:1, 34:1 and 36:1 to be the most abundant in yeast cells and used these in our analyses. The MRM mass data for each of these six lipid species is listed below

| Lipid Species | Parent Ion Mass | Daughter Ion Mass |
| --- | --- | --- |
| 32:1 PIP | 931.5 | 549.5 |
| 34:1 PIP | 959.5 | 577.5 |
| 36:1 PIP | 987.5 | 605.5 |
| 32:1 PIP<sub>2</sub> | 1039.5 | 549.5 |
| 34:1 PIP<sub>2</sub> | 1067.5 | 577.5 |
| 36:1 PIP<sub>2</sub> | 1095.5 | 605.5 |

The Sciex MultiQuant software was used to quantify the area under the peaks. For each sample, we analyzed the ratio of corresponding PIP_2_ and PIP lipid species by dividing the area under curve calculated for each lipid species of PIP_2_ with that of PIP lipid species with the same total carbon-chain length and number of double bonds. This normalized for the variations in lipid extractions between samples.

### PIP4K enzyme activity assay

S2R+ cells (2 million cells) were cultured in Schneider’s insect medium (HiMedia IML003) supplemented with 10% Fetal Bovine Serum and with antibiotics Penicillin and Streptomycin. Cells were transfected using Effectene (Qiagen, Cat no. 301425) based on the manufacturer’s protocol. PIP4K enzymes tagged to GFP were cloned in pUAST-attB plasmid and used for transfection in S2R+ cells. 48 hours post transfection, cells were pelleted down and washed with 1X PBS twice. The cells were lysed using 120ul of lysis buffer (50mM Tris/Cl-pH 7.5, 150mM NaCl, 4% protease inhibitor cocktail (PIC)). A medical syringe was used to mechanically lyse the cells. The resultant cell lysate was stored at −20 ^ͦ^ C. The cell lysates were further used for kinase assay.

### Bradford Assay and Western Blot

Bradford assay was used to measure the total amount of protein present in the cell lysate. Samples were prepared by adding 6X protein loading dye and heating them at 95 ^ͦ^C for 3 minutes. The proteins were separated using SDS-PAGE (10%). The proteins separated were transferred onto a nitrocellulose membrane using wet transfer protocol. The transfer was run at 100 volts for one and half hours following which the blot was rinsed and blocked with 10% BLOTO in 0.1 % TBST [20mM Tris/Cl – pH 7.6, 150mM NaCl, 0.1% Tween]. The blot was incubated overnight with primary antibody, anti-GFP (sc-9996) 1: 2,000 dilution or anti-GST (26H1) Mouse mAb#2624 1:3000 dilutions in 5% BSA in TBST. The membrane was then washed thrice in TBST and incubated with the secondary antibody, goat alpha mouse IgG1: 10,000 dilutions in 5% BSA in TBST for 3 hours. The blot was thoroughly washed and the proteins were visualised using ECL reagent (Bio-Rad) in an ImageQuant LAS 4000. Protein levels were normalised with tubulin levels to load equal quantities of protein for all samples for downstream assays.

### Mass spectrometry based kinase assay

600 picomoles of PI5P (phophatidylinositol-5-phosphate, Catalogue No. P5016-Echeleon biosciences) was mixed with 20 μl of 0.5 (mM) of Phosphatidylserine (PS) (P5660, Sigma). 100 ul of chloroform was added to this solution and dried in a centrifugal vacuum concentrator. To this mixture, 50 μl 10 mM Tris-HCl pH 7.4 was added and the mixture was sonicated for 3 minutes in a bath sonicator to form lipid micelles. 50 ml of 2X kinase assay buffer (100 mM Tris pH 7.4, 20 mM MgCl_2_, 140 mM KCl, and 2 mM EGTA containing 80 uM of cold ATP (10519979001, Roche) was added to each reaction tube. The mixture was kept on ice for 15 minutes. 1 μg equivalent of purified enzyme GST-hPIP4K2A was used as control for the experiment. 10 µg equivalent of protein was added from S2 cell lysate as enzyme source. The reaction was allowed to take place in an incubator at 30^0^C at 1000 rpm for 30 minutes. Reactions were stopped by adding 125 μl, 2.4 N HCl, 250 μl methanol and 250 μl chloroform. The mixture was vortexed vigorously and spun down for 5 minutes at 1000 xg to obtain clean phase separation. The lower organic phase was washed with equal volume of LPWS and vortexed and phase separated. The final organic phase obtained was processed further for chemical derivatization (for LC-MS/MS detection) to analyse the products of the reactions. Samples were run on a hybrid triple quadrupole mass spectrometer (Sciex 6500 Q-Trap) connected to a Waters Acquity UPLC I class system. Separation was performed either on a ACQUITY UPLC Protein BEH C4, 300Å, 1.7 μm, 1 mm X 100 mm column (186005590) or a 1 mm X 50 mm column (186005589), using a 45% to 100% acetonitrile in water (with 0.1% formic acid) gradient over 10 minutes or 4 minutes^21^.

### *In Vitro* kinase assay (Thin layer chromatography)

S2R+ cells were harvested by centrifugation at 1000 x g for 10 minutes at 4°C. The cell pellet was washed twice with ice-cold PBS. Cells were then homogenized in lysis buffer containing 50 mM Tris-Cl (pH 7.5), 1 mM EDTA, 1 mM EGTA, 1% Triton X-100, 50 mM NaF, 0.27 M sucrose, and 0.1% β-mercaptoethanol supplemented with freshly added protease and phosphatase inhibitors (Roche). The lysate was clarified by centrifugation at 1000 x g for 15 minutes at 4°C. Protein concentration in the lysate was determined using the Bradford assay according to the manufacturer’s instructions. Vacuum-dried PI5P substrate (final concentration 6 mM) and 20 mM phosphatidylserine were resuspended in 10 mM Tris (pH 7.4). Micelles were formed by sonication in a bath sonicator for 2 minutes. 50 μl of 2x PI5P4-kinase reaction buffer (containing 100 mM Tris pH 7.4, 20 mM MgCl_2_, 140 mM KCl, and 2 mM EGTA) supplemented with 20 mM ATP, 5 μCi [γ-³²P] ATP, and 10 μg of total cell lysate protein was added to the pre-sonicated micelles. The reaction mixture was incubated for 16 hours at 30°C. Lipids were extracted from the reaction mixture and resolved by one-dimensional TLC using a solvent system of chloroform: methanol: water: 25% ammonia (45:35:8:2 v/v/v/v). The resolved lipids were visualized using a phosphorimager.

### Generation of transgenic lines

The transgenic fly lines containing AqPIP4K, HvPIP4K and MbPIP4K on chromosome II were generated using site-specific φC31 integrase system^22^. These strains were used to generate fly strains which express AqPIP4K, HvPIP4K and MbPIP4K under the Gal4 promoter in the endogenous PIP4K knock-out flies.

### Larval salivary gland cell size measurement

The wandering third instar larvae were collected from the crosses mentioned (additional file) and salivary glands were dissected. A minimum of 7-8 salivary glands were collected per genotype. The glands were collected in 1X PBS. The glands were then fixed using 4% PFA solution for 15-20 minutes at 4^0^C. The glands were then washed with 1X PBS and kept in the same buffer until staining. The glands were incubated in Bodipy-F6-SM (membrane binding fluorescent dye) stain for 3 hours at room temperature. The glands were then stained with DAPI (1:2000) for 10 minutes at room temperature. The staining is done under dark conditions. They were then washed with 1X PBS 3 times, 10 minutes each. The salivary glands were then mounted on glass slides using 70% glycerol and imaged using confocal microscope (Olympus FV3000) at 10X magnification. The images obtained from confocal microscope were processed using ImageJ analysis tool. The total volume of the gland and number of nuclei were measured using VOLOCITY image analysis software. The average cell size was then calculated using the volume and number of nuclei. Significance in difference between the cell size of different genotypes was established using two tailed t-test in graph pad prism (Version 9.2).

### Body weight measurement of *D. melanogaster* larvae

Larvae of the required genotypes were generated by standard genetic crosses. Flies were allowed to lay eggs for 4 hours. The age of the larvae was calculated from the mid-point of the four-hour period. 25-30 newly hatched first instar larvae were transferred to new cut vials containing media and reared at 25 ^°^C. At time-points of 24, 72 and 96 hours post egg laying, larvae were collected and body weight was measured. Samples with ten larvae at a time were measured using precision balance. All body weight measurements were done for 100 larvae per genotype.

## Results

### Genes encoding PIP4K are present only in metazoan genomes

The PIP4K sequence from human (PIP4K2A) was used as query to search for homologs in the proteomes in UniprotKB and NR database of NCBI. The UniprotKB includes 2225 eukaryotes (of which 58 are unicellular eukaryotes), 8862 bacteria, 11478 viruses and 360 archaea proteomes. The NCBI dataset includes all protein sequences available to date. 250 homologs of PIP4K sequence were obtained from UniprotKB while 181 unique homologs were obtained from NCBI search, all of which contained the characteristic sequence features of PIP4K. A taxonomy assignment was done using the sequence IDs of the homologs and it was found that all PIP4K sequences belonged to organisms from metazoan kingdom (**Figure 1B**). In addition, PIP4K query sequences were also used to search proteomes of marine unicellular organisms (http://tara-oceans.mio.osupytheas.fr/ocean-gene-atlas/) that contained 371 proteomes. Overall, we found that of the 23296 proteomes surveyed, PIP4K sequence homologs were found only in metazoan genomes. They were not found in prokaryotes, marine unicellular organisms (http://tara-oceans.mio.osupytheas.fr/ocean-gene-atlas/) or unicellular eukaryotes.

As an additional step, PI3K, PI4K, PIP5K and PIP4K sequences were used as query to search the selected proteomes mentioned in Supplementary Table 2. Based on the presence of sequence motifs and conserved residues present in the activation loop, PIP4K sequences were differentiated from other phosphatidylinositol kinases. All the selected proteomes contained an intact PIP5K protein in them (Table 1). A phylogenetic tree of all PIP4K sequences (**Figure 1C**) revealed three distinct clusters based on bootstrap values: one that contained the sequences from early metazoans like Sponges, Cnidaria and Nematodes. The second cluster contained sequences from Arthropods and Molluscs and the third contained sequences from vertebrate proteomes. This third cluster could be further sub-divided into three sub-clusters corresponding to the three isoforms of PIP4K present in vertebrates. The similarity between the enzyme sequence was higher within each cluster compared to across clusters as inferred using the bootstrap values at each node. Thus, PIP4K sequences remain conserved across the metazoan tree of life starting from the earliest examples of metazoans through to mammalian genomes.

One representative organism, with studies related to PIP4K available in the literature, was selected from each clade in the phylogeny. A phylogenetic tree was generated using PIP4K sequence selected as mentioned above from each phylum to represent the evolution of PIP4K in metazoans (**Figure 1D**). It was observed that the PIP4K from earliest evolved metazoans form the base cluster and they are similar to PIP4K from invertebrates. The vertebrate PIP4K form a separate cluster and are distinct from these two groups of PIP4K sequences.

We then mapped the known conserved motifs and domains onto the selected PIP4K sequences (**Figure 1E**). The sequence identity between PIP4K from first evolved metazoans and those from both invertebrate and vertebrate proteins was around 40% while the sequence identity between invertebrate and vertebrate PIP4K was around 50-60%; the identity within the vertebrate clade was above 75%. All the sequences contained the catalytic PIPK domain (Pfam ID: PS51455). Two known motifs “DLKGS” and “MDYSLL” were also conserved in these sequences (**Figure 1F**). While the former is known to be important for substrate and ATP binding, the latter is important for ATP binding. Interestingly we found that an N-terminal region known to be important for substrate binding was highly variable which might suggest varied levels of enzyme-substrate interaction leading to variation in enzyme activity across PIP4K from different organisms^23^. However, the activation loop sequence remained highly conserved which strongly suggests the substrate specificity of these enzymes might be very similar (**Figure 1F**).

### Functional expression of ancient PIP4K

We transfected *AqPIP4K::GFP* into cultured *Drosophila* S2R+ cells. Western blots of these transfected S2R+ cell lysates revealed the expression of a protein of the expected M_r_ (**Figure 2B**). We also determined the subcellular distribution of AqPIP4K::GFP and found that it localizes to the plasma membrane and cytoplasm but is not localized in the nucleus (**Figure 2C**), a distribution similar to that of *dPIP4K::GFP* expressed in S2R+ cells. We also generated a kinase-dead AqPIP4K enzyme wherein the Asp residue of the MDYSLL motif was mutated to Ala (D274A) (**Figure 2A**). The localization and expression of the kinase-dead enzyme when expressed in S2R+ cells were found to be similar to that of the wild type protein (**Figure 2C**). The localization and protein expression of PIP4K from choanoflagellates (MbPIP4K) and Cnidaria (HvPIP4K) were also checked in S2 cells. Both MbPIP4K::GFP and HvPIP4K::GFP enzymes are expressed in S2 cells as shown by Western blots (**Figure 2B**). HvPIP4K localizes similar to the other metazoan PIP4K at the plasma membrane and cytoplasm in S2 cells. However, interestingly, MbPIP4K::GFP localizes only at the plasma membrane in S2 cells (**Figure 2C**). Since the localization of both wild-type and kinase-dead AqPIP4K enzyme remains same, any difference seen in the function of the two proteins can likely be attributed to the kinase activity of the protein.

**Figure 2:**
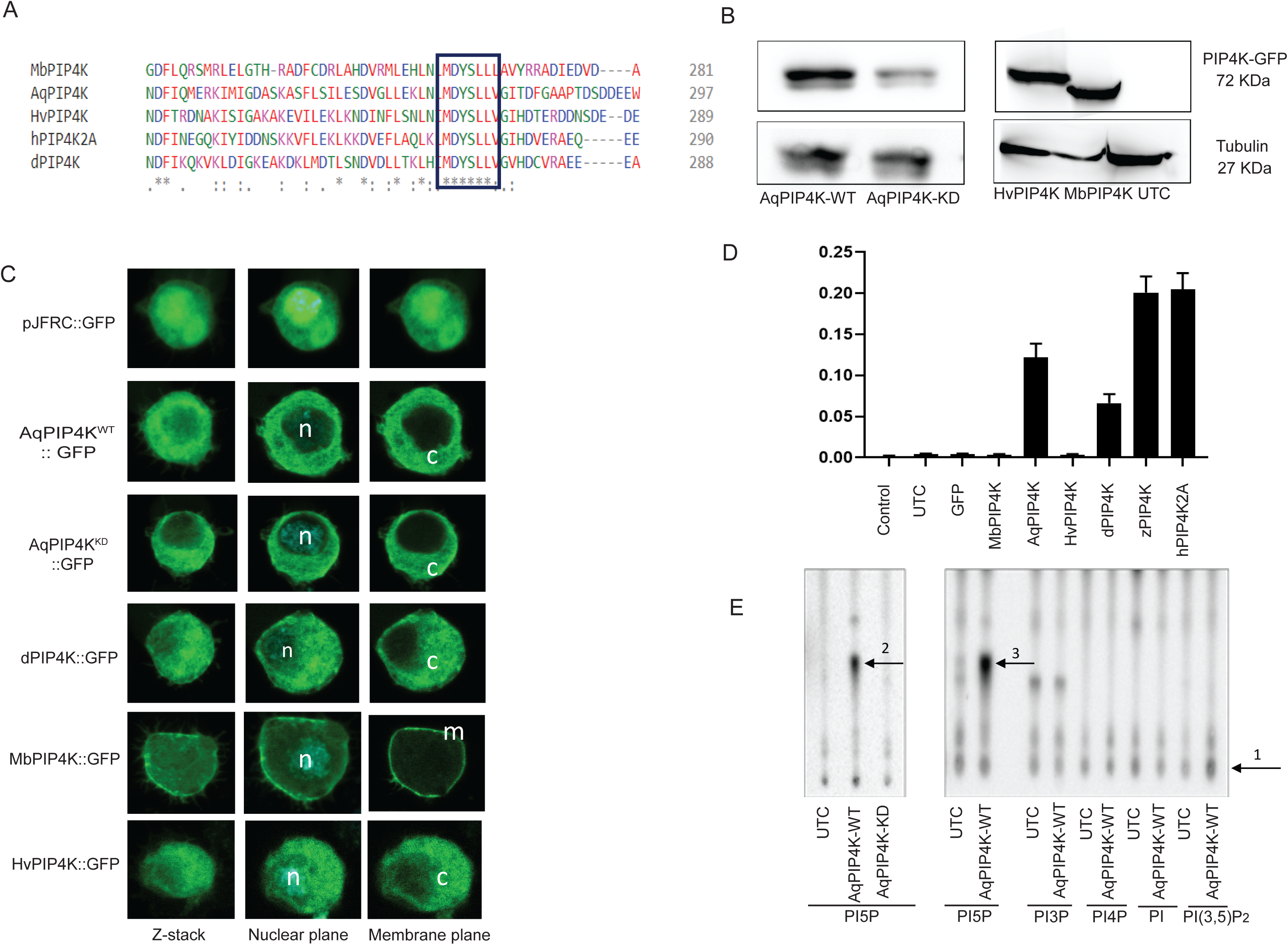
**(A)** Alignment of selected PIP4K sequence to show the generation of kinase-dead AqPIP4K enzyme by site directed mutagenesis. The aspartate residue of the “MDYSLL” motif has been mutated to Alanine (D275A). **(B)** Expression of wild-type AqPIP4K, MbPIP4K and HvPIP4K and kinase-dead AqPIP4K enzymes after transfection in S2 cells, Tubulin expression has been used as internal control (western blot). **(C)** Localization of wild-type AqPIP4K, MbPIP4K and HvPIP4K and kinase-dead AqPIP4K enzyme tagged to GFP in S2 cells as observed by confocal microscopy (10X, GFP channel for protein). Vector expressing only GFP was used as control. Localization of *dPIP4K::GFP* was used for comparison. **(D)** Mass spectrometry based kinase assay to check the enzyme activity of selected PIP4K enzymes. The graph represents the response ratio (PIP_2_/PIP) for the *in-vitro* kinase assay for each of the selected PIP4K enzyme. (D,i) Lipid kinase activity of wild-type and kinase dead mutant AqPIP4K. TLC showing the formation of radiolabelled PI(4,5)P_2_ from PI5P by while AqPIP4K-Kd does not. UTC-untransfected control reaction **(E)** Lipid substrate utilization by wild-type AqPIP4K. TLC showing radiolabelled products formed from Phosphatidylinositol 5-phosphate (PI5P), phosphatidylinositol 3-phosphate (PI3P), phosphatidylinositol 4-phosphate (PI4P), phosphatidylinositol (PI), and phosphatidylinositol 3,5-bisphosphate (PI(3,5)P_2_). Arrow head indicates the substrate and formation of a product. 1-Substrates, 2 and 3 – Product PI(4,5)P_2_.

### AqPIP4K shows enzymatic activity *in vitro*

We tested the enzymatic activity of AqPIP4K using an *in vitro* biochemical assay. Lysates of S2R+ cells transfected with a PIP4K gene were incubated with its best-known substrate PI5P and ATP. The consumption of PI5P and the formation of the product PI(4,5)P_2_ was monitored using mass spectrometry. As a positive control, as previously reported ^21^, we monitored the formation of PI(4,5)P_2_ from PI5P by hPIP4K, zPIP4K and dPIP4K (**Figure 2D**). Under these conditions we found that transfection of S2R+ cells with AqPIP4K resulted in the robust conversion of PI5P to PI(4,5)P_2_. Mass spectrometry based kinase assay was performed to test the kinase activity of the PIP4K from *M. breviolis, and H. vulgaris*. Under these conditions, MbPIP4K and HvPIP4K were not active and only AqPIP4K showed conversion of PI5P to PIP_2_ in the *in vitro* kinase assay. This indicates that AqPIP4K is active and might be functionally important in *A. queenslandica.* (**Figure 2D**).

### Conserved substrate specificity between AqPIP4K and PIP4K from complex metazoans

*In vitro* and *in vivo* studies of PIP4K from higher metazoans had demonstrated a high degree of substrate specificity for the enzyme. PIP4K can utilize PI5P as a substrate but not PI4P ^9,18^. Further, it can use PI3P as a substrate with low efficiency ^4,21^. To determine the substrate specificity of AqPIP4K, we used a kinase assay where ^32P^γATP is used to detect radiolabelled lipid products separated on a TLC. Under these consitions, AqPIP4K was able to convert PI5P into radiolabelled PI(4,5)P_2_ whereas a non-ATP binding kinase dead mutant (D273A) was unable to do so (**Figure 2E**). Under equivalent conditions AqPIP4K could not phosphorylate PI4P and was able to convert PI3P into PI with very low efficiency (**Figure 2E**). AqPIP4K was also unable to phosphaorylate phosphatidylinositol (PI) and phosphatidylinositol-3,5-bisphosphate (PI(3,5)P_2_).

### AqPIP4K has a unique and non-redundant biochemical activity *in vivo*

To test if the ability of AqPIP4K to specifically phosphorylate PI5P but not PI4P *in vitro* is also conserved *in vivo*, we used the *S. cerevisiae* model.The *S. cerevisiae* genome encodes a single gene *mss4*, that has previously been shown to encode a PIP5K (**Figure 1A**). Deletion of *mss4* is lethal in *S. cerevisiae*, therefore, SD102 carrying the temperature sensitive allele *mss4-2^ts^* shows wild type growth at permissive temperature (28 or 30°C) but shows almost no growth at the restrictive temperature (38°C) (**Figure 3A i and ii)**. Previous studies have shown that this growth defect of *mss4-2^ts^* can be rescued by reconstituting with hPIP5K but not hPIP4K^24^ and therefore is an excellent model to test the function of a new PIPK and classify it either as a PIP5K or a PIP4K. To test whether AqPIP4K is a PIP5K or PIP4K *in vivo*, we reconstituted *mss4-2^ts^* with this gene. Cells containing PYES2-AqPIP4K, PYES2-hPIP4K2A and PYES2-MSS4::GFP were allowed to grow at permissive and restrictive temperatures to the early logarithmic phase. The functional activity of any reconstituted PIPK was measured by its ability to restore the growth defect in *mss4-2^ts^* at the restrictive temperature. Under our experimental conditions, MSS4::GFP was able to restore growth (**Figure 3A iii**) and we found that as previously reported ^24^, hPIP5K was able to rescue the growth defect of *mss4-2^ts^* at 38^0^C; however, hPIP4K2A was unable to do so (**Figure 3A iii, Figure 3B i**). Under the same conditions, AqPIP4K was unable to restore normal growth in *mss4-2^ts^* (**Figure 3A iv, Supplementary Figure 1F**)). Under the same conditions, MbPIP4K, HvPIP4K and dPIP4K were also not able to restore growth in *mss4-2^ts^* (**Figure 3B iii, iv**). The number of colonies formed in each of the above mentioned cases has been quantified as represented in **Figure 3C**. These findings indicate that AqPIP4K cannot function as a PIP5K *in vivo*.

**Figure 3:**
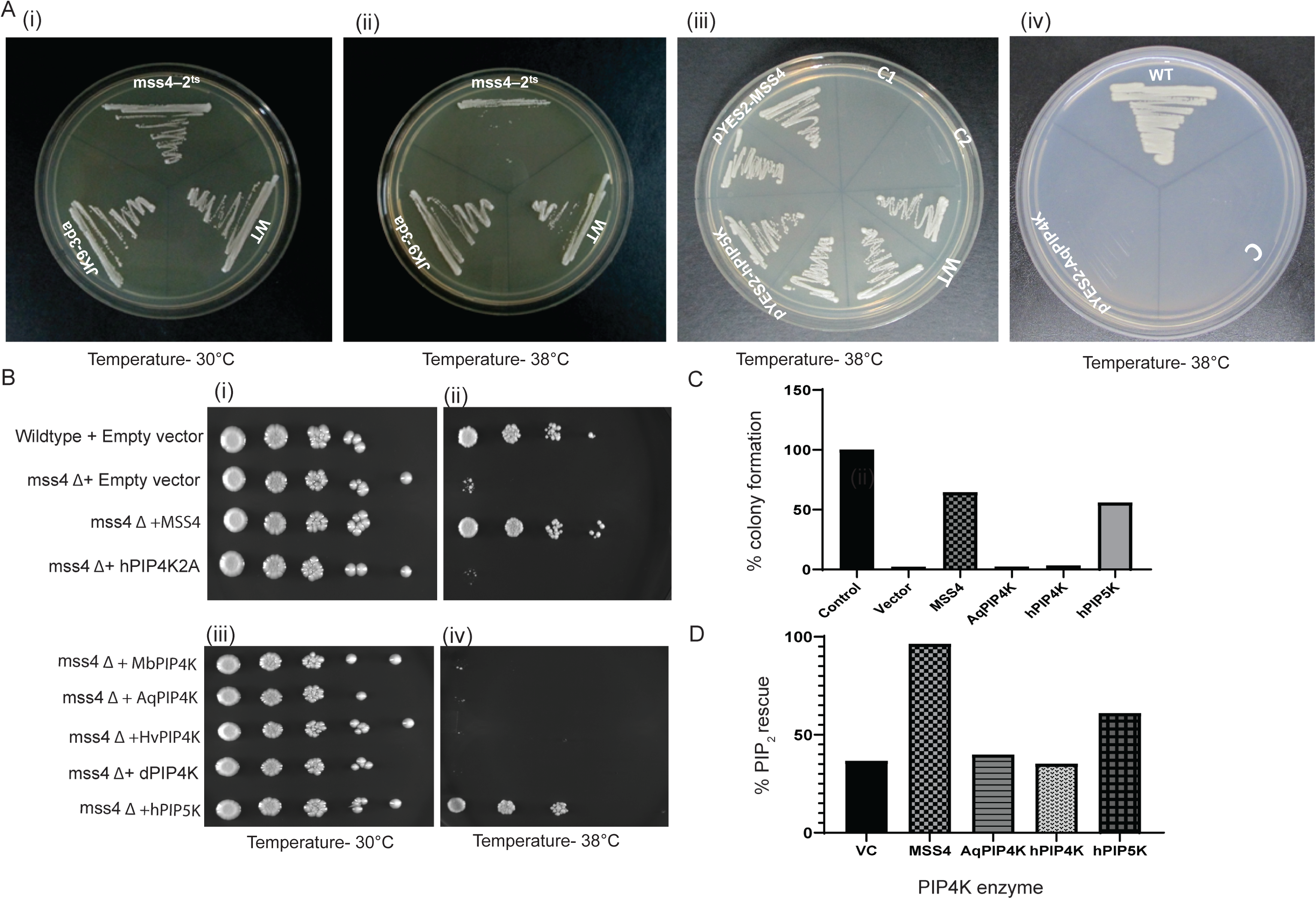
**(A) (i-ii)** Growth of yeast cells, wild-type (JK9–3d, background strain), MH-272 (JK9–3da/αhis3Δ/his3Δ HIS4/HIS4) and temperature sensitive strain (SS102^ts^, MH272–1da/αmss4::HIS3MX6/MSS4-2^ts^) at 30 ^ͦ^C and 38 ^ͦ^C. **(iii and iv)** Growth of temperature sensitive yeast cells reconstituted with MSS4, hPIP5K1A and AqPIP4K at 38 ^ͦ^C. C, C1-untransformed control and C2-empty vector **(B) (i-ii)** Growth of yeast cells for colony counting at 30 ^Cͦ^ and 38 ^Cͦ^. The cells plated are wild-type, mss4 Δ, *mss4Δ* reconstituted with MSS4 and *mss4Δ* reconstituted with hPIP4K2A. **(iii and iv)** Growth of yeast cells for colony counting at 30 ^Cͦ^ and 38 ^ͦ^C. The cells plated are mss4 Δ reconstituted with MbPIP4K, *mss4Δ* reconstituted with AqPIP4K, *mss4Δ* reconstituted with HvPIP4K, *mss4Δ* reconstituted with dPIP4K and *mss4Δ* reconstituted with hPIP5K**.(C)** Percentage colony forming units quantified from Figure 3(B). **(D)** Quantification of levels of the product of PIP4K (PIP_2_) as percentage PIP_2_ rescue in yeast upon over expression of mss4, AqPIP4K, hPIP4K and hPIP5K

To correlate the ability to restore growth in *mss4-2^ts^* with the underlying biochemical defect, we measured the levels of PI(4,5P)_2_, the product of MSS4 activity. Complementation of *mss4-2^ts^* with *mss4* was able to elevate the level of PI(4,5)P_2_. Likewise complementation with hPIP5K was able to also elevate PI(4,5)P_2_ levels; however complementation with hPIP4K and AqPIP4K was unable to elevate PI(4,5)P_2_ levels (**Figure 3D**). These findings indicate that the inability of AqPIP4K to restore growth in *mss4-2^ts^* is correlated with its inability to elevate PI(4,5)P_2_ levels

### AqPIP4K is able to rescue growth phenotypes of dPIP4K mutants

Since AqPIP4K showed a biochemical activity similar to dPIP4K, zPIP4K and hPIP4K2A, we wondered if the *Amphimedon* enzyme would be able to complement for the function of a metazoan PIP4K. For this we used the *Drosophila* model; in flies, loss of *dPIP4K* results in a delay in development, a growth deficit and reduced salivary gland cell size^12^. We tested the effect of reconstituting AqPIP4K in *dPIP4K* null mutants (*dPIP4K*^29^)^12^. When reconstituted throughout fly tissues (using Act>Gal4), *AqPIP4K* was able to largely rescue the reduced body weight of *dPIP4K*^29^ (**Figure 4A**) at 72 hours post eclosion and also at other time points throughout larval development **(Supplementary Figure 1A, 1B, 1C**); as a positive control we reconstituted *dPIP4K*^29^ with *dPIP4K* which was also able to rescue the growth defect to a similar extent as *AqPIP4K* (**Figure 4B**). The size of salivary gland cells in the 3^rd^ instar larvae of *dPIP4K*^29^ is known to be reduced (**Figure 4C**); we tested in if AqPIP4K was able to rescue this reduced cell size and found that like *dPIP4K*, *AqPIP4K* was able to rescue cell size in *dPIP4K*^29^ (**Figure 4C, 4D, 4E**). In order to check whether kinase activity of the AqPIP4K enzyme is important for its function, we mutated the Asp residue of the ‘MDYSLL’ motif of AqPIP4K to Ala residue (D275A). This mutation has shown to render the protein kinase-dead in mammalian and invertebrate systems^25^. The rescue of both the larval growth defect as well as salivary gland cell size was dependent on kinase activity as a kinase dead version of AqPIP4K was unable to rescue either of these phenotypes (**Figure 4A, 4D**). PIP4K from choanoflagellates (MbPIP4K) and Cnidaria (HvPIP4K) however, when reconstituted in the null mutant flies was unable to rescue the salivary gland cell size in the same setting (**Supplementary Figure 1D, 1E**).

**Figure 4:**
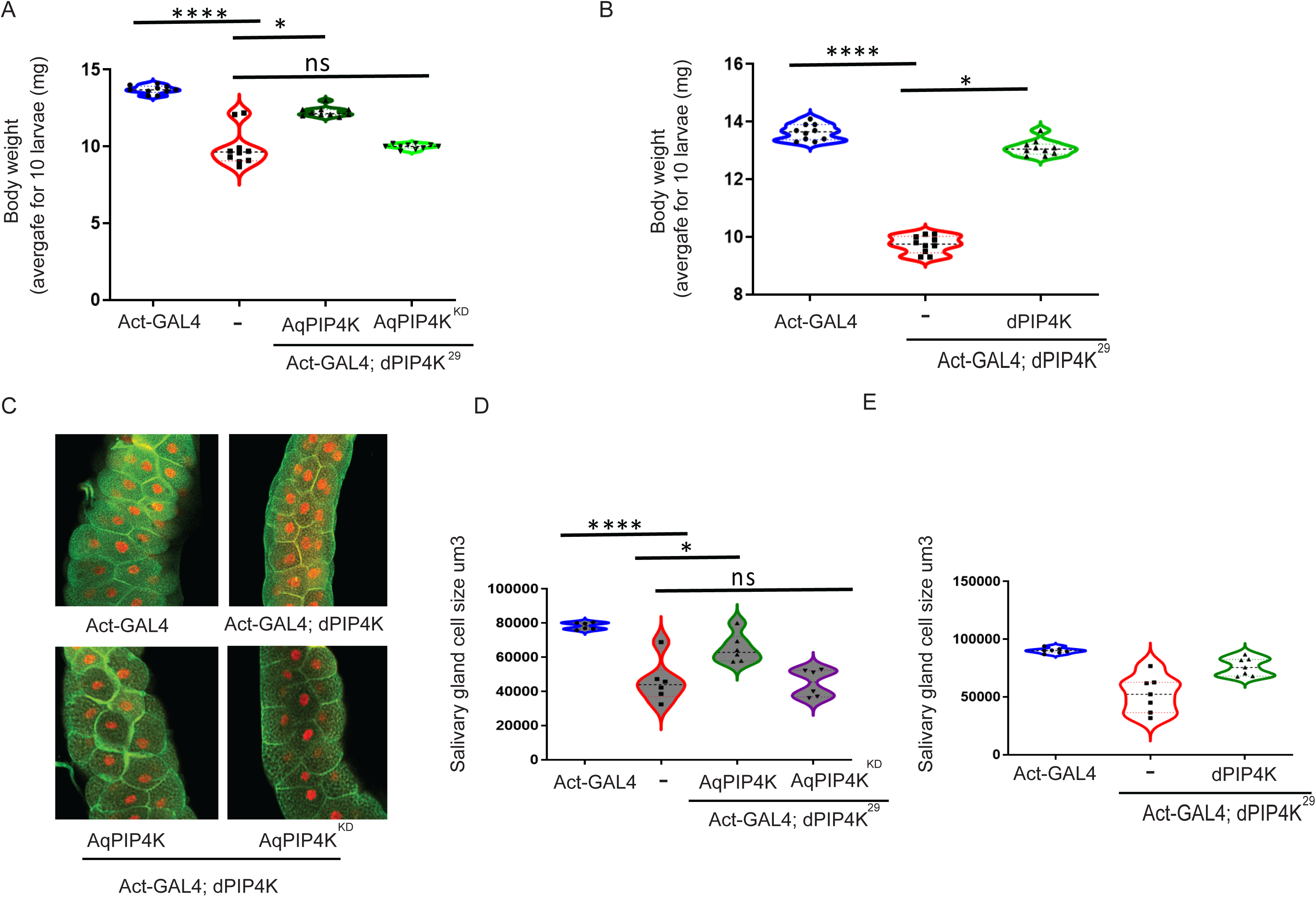
**(A)** Body weight of larvae measured at 72 hours post egg laying. Each point on the graph represents average weight of 10 larvae (total of 100 larvae measured per genotype). The genotypes include wild-type, *dPIP4K*^29^, *dPIP4K*^29^ larvae reconstituted with AqPIP4K and AqPIP4K-KD. **(B)** Body weight of larvae measured at 72 hours post egg laying. Each point on the graph represents average weight of 10 larvae (total of 100 larvae measured per genotype). The genotypes include wild-type, *dPIP4K*^29^*, dPIP4K*^29^ larvae reconstituted with dPIP4K. **(C)** Representative confocal images of salivary glands from the genotypes *Act-Gal4/+, Act-Gal4/+;; dPIP4K*^29^, *Act-Gal4>AqPIP4K;;dPIP4K*^29^, *Act-Gal4>AqPIP4K-KD;; dPIP4K*^29^. Cell body is marked green by BODIPY conjugated lipid dye, and the nucleus is marked by DAPI shown in red. Scale bar indicated at 100 μm. **(D)** Salivary gland cell size measurements from third instar wandering larvae wild-type (*w^11^*^18^), *dPIP4K*^29^ (flies with no expression for dPIP4K protein), *dPIP4K*^29^ larvae reconstituted with AqPIP4K and AqPIP4K^KD^ respectively. **(E)** Salivary gland cell size measurements from third instar wandering larvae wild-type (*w^11^*^18^), *dPIP4K*^29^ (flies with no expression for dPIP4K protein), *dPIP4K*^29^ larvae reconstituted with dPIP4K.

## Discussion

The use of PI as signal transduction elements is a conserved features of eukaryotes. The phosphorylated derivatives of PI such as PI3P, PI(4,5)P_2_ and PI(3,5)P_2_ are used to control core sub-cellular processes such as vesicular transport and cytoskeletal re-organization, processes that are conserved in all eukaryotes. These lipids have been isolated from unicellular eukaryotes through to metazoans and the lipid kinases and phosphatases that control their levels are also found in all genomes of eukaryota. However, some phosphoinositides such as PIP_3_ have been reported only in metazoans and the lipid kinases and phosphatases that regulate its levels are also found only in metazoan genomes **(Supplementary Figure 2B**). In metazoan biology, communication and coordination across cells is critical and this is mediated through chemical signals, growth factors such as insulin. Receptors for many such growth factors, receptor tyrosine kinases^7,26,27^, are found uniquely in metazoan genomes and mediate signal transduction by triggering the activity of PI3KC1 to generate PIP_3_. Thus, all three core elements of hormone signalling in multi-cellular organisms, namely the ligand (insulin), its receptor (receptor tyrosine kinase) and PI3KC1 are found only in metazoans **(Supplementary Figure 2A**). Given this observation, one might also predict that regulators of PI3KC1 might also have evolved in metazoans. PI3KC1 is regulated by multiple processes ^8^; several regulators such as Ras and G_βγ_ are found in all eukaryota including both unicellular and multicellular organisms although they operate in contexts where GPCRs activate PI3KC. In this study, we found that a recently described regulator of PI3KC1, PIP4K is found only in the genomes of metazoans. Deletion of each of the three mammalian PIP4K genes in mouse models had revealed phenotypes consistent with enhanced receptor tyrosine kinase mediated signalling; PIP4K2A-epidermal growth factor signalling; PIP4K2B-insulin receptor signalling and PIP4K2C-T-cell receptor signalling ^14,28^. Together with recent studies in *Drosophila* ^13^ and human cells ^14^ that have shown PIP4K as a key biochemical regulator of PI3KC1 signalling, these findings indicate that PIP4K is a key regulator of receptor tyrosine kinase signalling. Remarkably genes encoding PIP4K are found in the earliest examples of metazoa, namely the coral *A.queenslandica* and in choanoflagellates, organisms that can exist transiently as groups of cells, i.e multicellular. What is the function of PIP4K in such ancient metazoans? Analysis of biochemical activity *in vitro* showed that under equivalent conditions both the *Amphimedon* and human enzyme, although separated by 50Mya years of evolution ^29^ have equivalent biochemical activity, namely the ability to specifically convert PI5P into PI(4,5)P_2_. Further, *in vivo* analysis using the yeast model showed that AqPIP4K also demonstrates the exquisite substrate specificity of the human enzyme being unable to convert PI4P into PI(4,5)P_2_. The *Amphimedon* genome also encodes an independent gene that is likely to function as a PIP5K. These indicate that the earliest known metazoan genome encodes two distinct enzymes to generate PI(4,5)P_2_, a PIP5K [PI4P to PI(4,5)P_2_] and a PIP4K [PI5P to PI(4,5)P_2_]. This observation suggests that the specific biochemical activity of PIP4K is required to support an unique aspect of metazoan biology in *Amphimedon*. What might this process be? We suggest that co-ordinated communication between thcells is the most likely function; such communication is typically mediated by hormone receptor interactions coupling to intracellular phosphoinositide signalling The identity of the hormone and receptor tyrosine kinase that works in this setting is not clear. Analysis of the *Amphimedon* genome^30^ reveals the presence of ca.150 genes encoding receptor tyrosine kinase like proteins with many having unique extracellular domains compared to their counterparts in eumetazoans; a direct homolog of the insulin receptor could not be identified. Thus, the identity of the specific RTK and the ligands that may bind to these receptors and couple to PI3KC1 remains to be determined. A gene encoding PI3KC1 is found in all metazoans including one in *A.queenslandica* (**Supplementary Figure 2C**) with the catalytic and other domains also conserved across proteins encoded in metazoans Thus, key elements of the signal transduction machinery required for PIP4K to regulate PI3KC1 in *Amphimedon* are encoded in the genome of this early metazoan. Although we could not experimentally test the biology regulated by PIP4K in *Amphimedon*, our analysis of AqPIP4K in *Drosophila* provides compelling evidence that AqPIP4K is able to regulate receptor tyrosine kinase activated PI3KC1 dependent processes in much the same way as shown for its eumetazoan counterparts ^13,14^.

How did PIP4K genes evolve in metazoans to regulating growth and developmentGiven that the related enzymes PIP5K is present in all eukaryotes, a plausible explanation is the PIP4K evolved from this ancestral PIP5K gene. Studies using sequence alignment and conservation between PIP4K and PIP5K suggest that gene duplication and neofunctionalization of PIP5K gene during metazoan evolution might have led to the emergence of PIP4K gene^31^. The ability if the new PIP4K to regulate PI3KC1 may then have positively selected the new gene. Further studies need to be performed to understand the distinctive and unique regulatory role (as compared to other regulators) of PIP4K on PI3KC1 activity during receptor tyrosine kinase signalling. There is uncertainty in the evolution of early metazoan. It is unclear whether the sponge-sister clade or ctenophore sister clade supported metazoan evolution as both these clades share unique features that could have shaped metazoan evolution^32^. There is a possibility that during evolution there have been some genes lost in sponges and some which have evolved independently in sponges and cnidaria^32^. These studies also suggest bilaterians and cnidaria have evolved by means of parallel evolution from sponges. In the context of other ancient PIP4K it is possible that since choanoflagellates exist as both single-cell and a transient multicellular state and do not have the characteristics of metazoans, PIP4K does not play any important functional role in these. This may explain the lack of functionality when MbPIP4K is expressed in *Drosophila* PIP4K mutants. It was however surprising that Cnidaria HvPIP4K was not active in either biochemical or functional assays. Based on theories of parallel evolution between cnidarians and sponges during early metazoan evolution, it is possible that the PIP4K gene was retained functional in one lineage and not in other^29 32^. Nonetheless, our findings position PIP4K as a key signal transduction enzyme in mediating cell-cell communication through hormones in the most ancient metazoans.

## Acknowledgements

This work was supported by the Department of Atomic Energy, Government of India, Project Identification No. RTI 4006 and Wellcome-DBT India Alliance Senior Fellowship to PR (IA/S/14/2/501540). RS is a J.C. Bose National Fellow (JBR/2021/000006) from the Science and Engineering Research Board, India. RS would also like to thank Bioinformatics Centre Grant funded by the Department of Biotechnology, India (BT/PR40187/BTIS/137/9/2021) and the Institute of Bioinformatics and Applied Biotechnology for the funding through her Mazumdar-Shaw Chair in Computational Biology (IBAB/MSCB/182/2022). We thank the Drosophila, mass spectrometry, Imaging and High-performance computing facility at NCBS for support.

---

**Supplementary Figure S1: (A, B and C)** Body weight of larvae measured at 42, 72 and 96 hours post egg laying respectively. The graph represents average weight of 10 larvae (total of 100 larvae measured per genotype). The genotypes include wild-type, *dPIP4K*^29^ and *dPIP4K*^29^ larvae reconstituted with AqPIP4K at 24hours, 72 hours and 96 hours post eclosion. **(D)** Salivary gland cell size measurements from third instar wandering larvae wild-type (*w^11^*^18^), *dPIP4K*^29^ (flies with no expression for dPIP4K protein), dPIP4K^29^ larvae reconstituted with MbPIP4K. **(E)** Salivary gland cell size measurements from third instar wandering larvae wild-type (*w^11^*^18^), *dPIP4K*^29^ (flies with no expression for dPIP4K protein), *dPIP4K*^29^ larvae reconstituted with HvPIP4K. **(F)** Western blot to show the expression of AqPIP4K in yeast transformed cells upon induction with galactose. VC-vector control, 1-Transformant 1, 2-Transformant 2, M-Marker and C-*Drosophila* expressed AqPIP4K as control. Actin was used as loading control.

**Supplementary Figure S2: (A)** Conservation of genes involved in the insulin signalling pathway in Human, *D.melanogaster* and *A.queenslandica*. Orange and blue squares indicate genes present in Human and *Drosophila g*enomes. Pink squares indicates orthologs found in *A.queenslandica*. Yellow squares indicate partial sequence found in *A.queenslandica.* **(B)** The genes in the PI signalling pathway which could have possible interaction with PIP4K in regulating its function. Genes marked in green are present in *A.queenslandica* genome while genes marked in red are absent. **(C)** Domains marked on PI3K proteins of Human, *D.melanogaster* and *A.queenslandica.* The image was obtained using the IBS server. The length of each domain is marked below the domains.

## Notes

### Competing Interest Statement

The authors have declared no competing interest.

