## Supplementary figures and images for "The conserved biochemical activity and function of an early metazoan phosphatidylinositol 5 phosphate 4-kinase regulates growth and development"

### supplementary figures

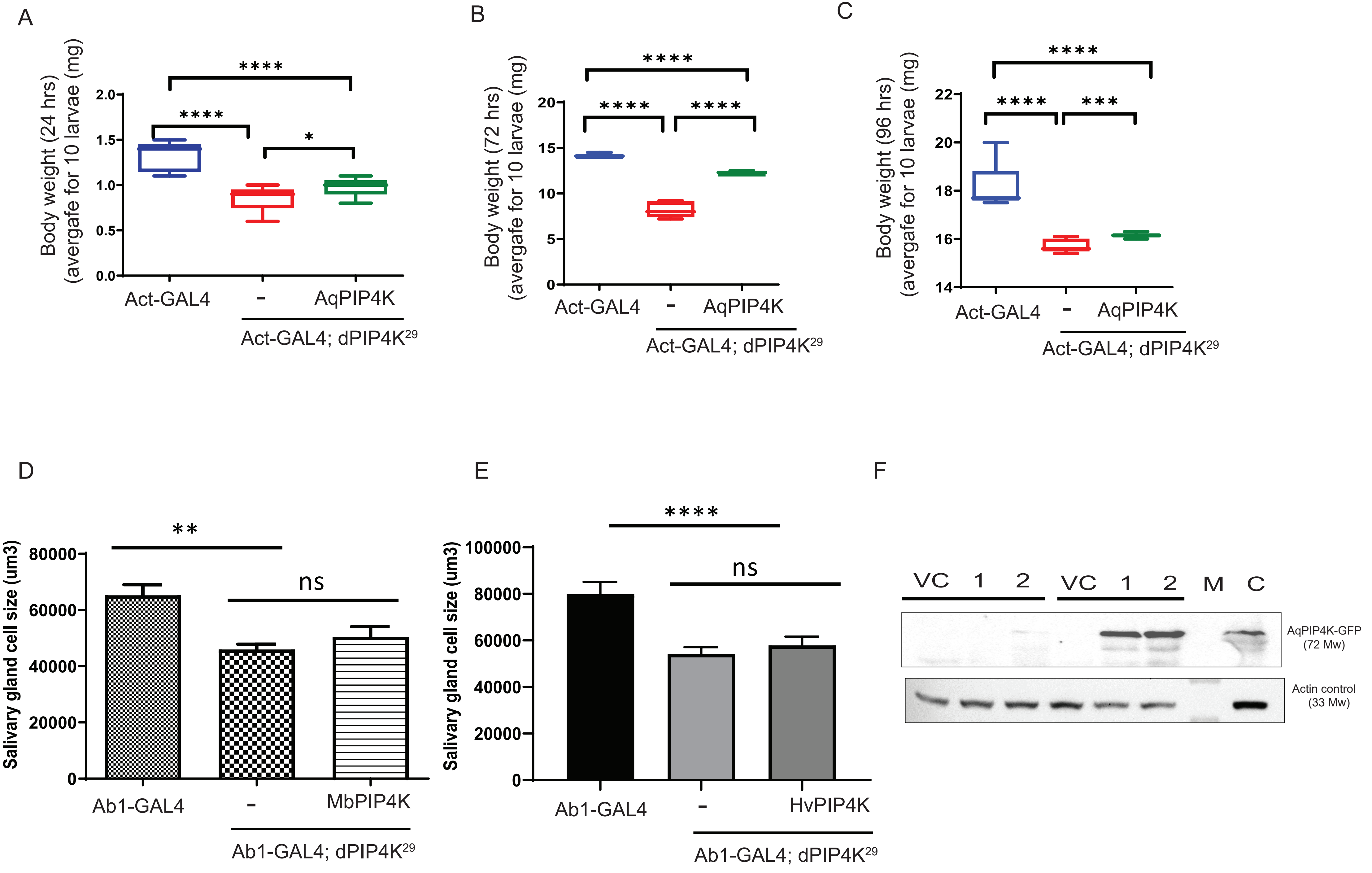

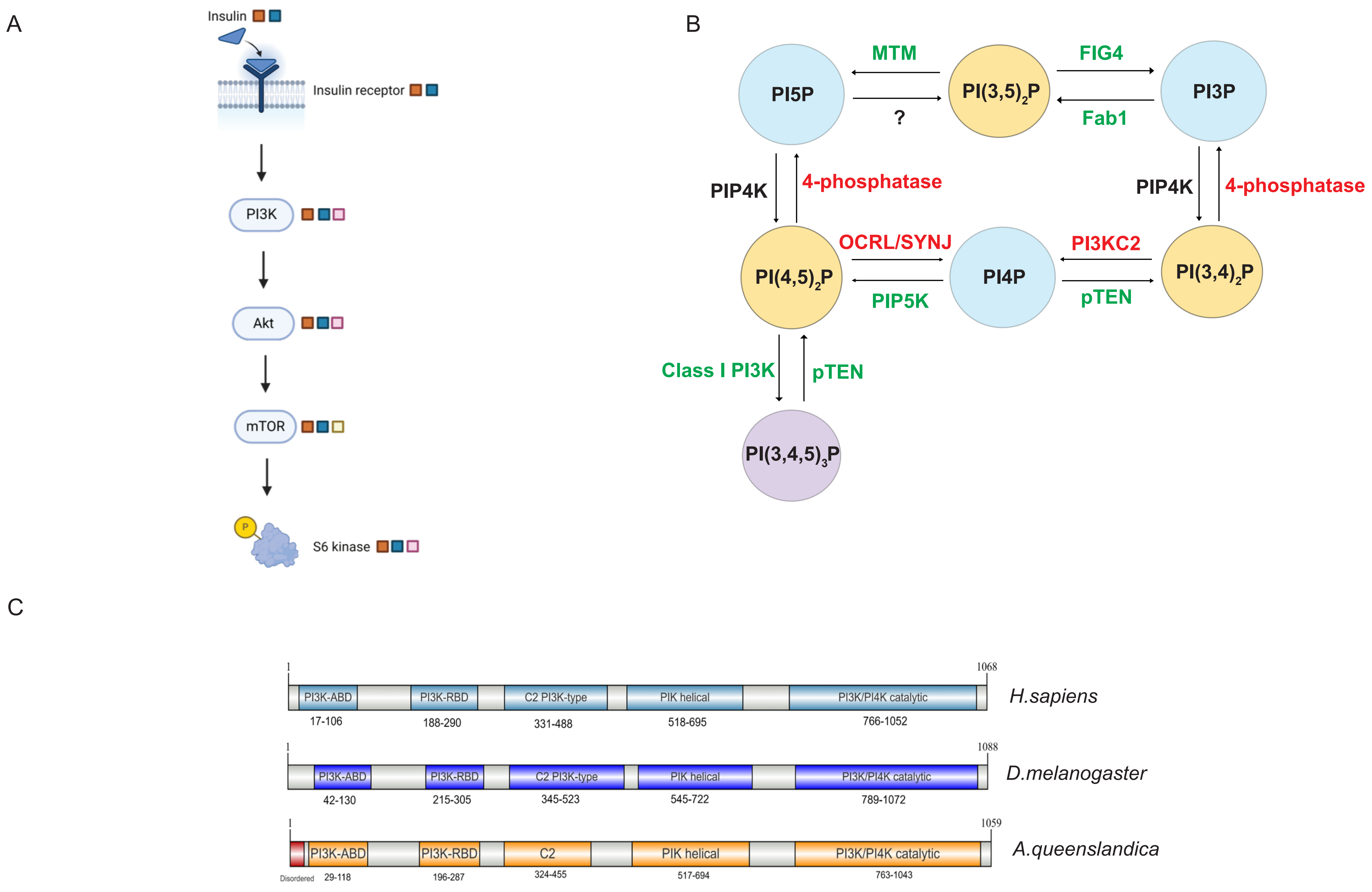


SupplementaryFigure_2_KH_Evolution
