## Supplementary Tables for "The conserved biochemical activity and function of an early metazoan phosphatidylinositol 5 phosphate 4-kinase regulates growth and development"

**Supplementary Table 1**: NCBI IDs of phosphatidylinositol kinases and phosphatidylinositol phosphate kinases selected as query to search for homologs in selected genomes.

| **Name of the protein** | **Refseq ID** |
|---|---|
| PI3K3 | NP_002638.2 |
| PI4P3KC2A | NP_002636.2 |
| PI4P3KC2B | NP_002637.3 |
| PI4P3KC2C | NP_004561.3 |
| PI45P3KA | NP_006209.2 |
| PI45P3KB | NP_006210.1 |
| PI45P3KD | NP_005017.3 |
| PI3P5K | NP_055855.2 |
| PI4KA | NP_060895.1 |
| PI4KB | NP_060793.2 |
| PI4P5K1A | NP_001129108.1 |
| PI4P5K1B | NP_003549.1 |
| PI4P5K1C | NP_001182662.1 |
| PI5P4K2A | NP_005019.2 |
| PI5P4K2B | NP_003550.1 |
| PI5P4K2G | NP_001139731.1 |

**Supplementary table 2**: List of multicellular organisms selected for genome wide search of PIP4Ks and PIP5Ks sequences

| **Name of the organism** | **Representative phylum** |
|---|---|
| *Dictyostelium* | Amoebozoa |
| *Arabidopsis* | Plantae |
| *Neurospora* | Fungi |
| *Saccharomyces* | Yeast |
| *Monosiga* | Holozoa |
| *Amphimedon* | Porifera |
| *Nematostella* | Cnidaria |
| *Helobdella* | Annelida |
| *Lottia* | Mollusca |
| *Drosophila* | Insecta |
| *Caenorhabiditis* | Nematoda |
| *Branchiostoma* | Chordata |
| *Homo sapiens* | Chordata |
